# Impact of chronic alcohol and stress on mid-life cognition and locus coeruleus integrity

**DOI:** 10.1101/2025.11.20.689500

**Authors:** O Revka, SJ Belculfine, L Fitts, KE Nippert, CAF Teves, PM Reis, S Tenney, BE Packer, I Garcia Alvarez, O Milstein, M Coutinho da Silva, DE Moorman, EM Vazey

**Affiliations:** Department of Biology, University of Massachusetts Amherst; Department of Psychological and Brain Sciences, University of Massachusetts Amherst; Neuroscience and Behavior Program, University of Massachusetts Amherst

**Keywords:** alcohol use disorder (AUD), reversal learning, cognitive flexibility, aging, dementia

## Abstract

**Background:** Excessive alcohol consumption and stress are associated with structural and functional alterations in the brain and impaired cognition. However, the persistence of long-term neural impacts after alcohol and stress are less understood. This study investigated midlife cognition and neuropathological changes following a history of alcohol and stress exposure.

**Methods:** C57BL/6J mice acclimated to ethanol drinking (15% v/v) before exposure to four cycles of alternating chronic intermittent ethanol (CIE) vapor exposure and repeated forced swim stress (FSS), with control groups exposed to air and no stress (AIR/NS). After three months of abstinence, mice were evaluated at midlife (11 months old) on volitional drinking and a final CIE/FSS challenge for stress induced drinking. Spatial learning and cognitive flexibility were assessed using the Barnes maze before brains were collected to evaluate locus coeruleus integrity at 12 months old.

**Results:** CIE/FSS increased volitional alcohol intake, and this drinking phenotype persisted through to midlife despite extended abstinence. CIE/FSS mice showed intact spatial learning but impaired flexibility in the Barnes maze reversal phase. Flexibility impairments were driven by decreased time in the target quadrant and increased errors during the reversal test compared to AIR/NS. Furthermore, CIE/FSS mice showed pathological measures of reduced locus coeruleus integrity common to dementia related disorders, including elevated markers of oxidative stress, apoptosis and reduced autoinhibitory function.

**Conclusions:** Our findings highlight the long-lasting impact of alcohol and stress exposure on cognition, with flexibility impairments persisting into midlife. In addition to cognitive changes, alcohol and stress history produced pathological changes in the locus coeruleus, an area known to mediate cognitive flexibility via its forebrain projections. Together, these results give an insight into the long-lasting impacts of chronic alcohol and stress and how they may accelerate age-related cognitive decline.

## Introduction

Alcohol use disorder (AUD) is characterized by compulsive alcohol consumption despite adverse consequences, and loss of control over drinking behaviors. It remains a significant cause of global mortality and morbidity, with major long term health consequences (Carvalho et al., 2019). Chronic alcohol consumption is associated with numerous negative effects on brain structure and function with heavy alcohol use facilitating age related declines in brain health. Alcohol history can accelerate brain aging including neuronal loss, reduced gray and white matter volume, reduced neurogenesis, and alterations in synaptic transmission and neuronal excitability (Avchalumov et al., 2021; Daviet et al., 2022; Funk-White et al., 2023; Mende, 2019; Nunes et al., 2019; Pleil et al., 2015; Smith et al., 2025; Sullivan and Pfefferbaum, 2023; Vetreno et al., 2011).

Stress is a critical contributor to AUD. Stress exposure alters activity in key brain regions involved in behavioral control and reward processing, promoting binge drinking and impairing control of alcohol consumption behaviors (Sinha, 2022). The relationships between alcohol and stress are bidirectional. Chronic alcohol impairs stress coping mechanisms by exacerbating anxiety and negative emotion regulation and disrupts neural responses in brain regions such as amygdala, dorsal striatum, prefrontal cortex, and orbitofrontal cortex (Bloch et al., 2022; den Hartog et al., 2020; Lu and Richardson, 2014; Rodberg et al., 2017). As a result, stress and alcohol intake act together in a positive feedback loop enhancing the negative effects of each other and increasing potential long-term damage.

Studying the impacts of stress in conjunction with models of AUD can identify specific synergistic vulnerabilities in brain physiology and function. Both chronic alcohol and stress have been implicated as risk factors for dementia (Livingston et al., 2024). However, the mechanisms linking chronic alcohol consumption, stress, and dementia remain poorly understood. Clarifying these interactions could reveal therapeutic targets for managing the long-term neurological risks associated with AUD.

One brain region sensitive to both alcohol and stress and heavily implicated in the pathophysiology of dementia is the nucleus locus coeruleus (LC) (Chan-Palay, 1991; Ehrenberg et al., 2023; Reyes, 2025; Vazey et al., 2018). LC is the brain’s primary source of norepinephrine (NE), projects extensively to key cortical regions involved in executive functions and plays a central role in attention, arousal, learning, memory, sensory processing, and stress regulation. These executive functions are impaired by chronic alcohol misuse and stress exposure. Additionally, LC is particularly vulnerable to neurotoxic impacts, making it a critical area for understanding the molecular mechanisms of alcohol- and stress-induced cognitive deficits.

To identify the consequences of excessive alcohol and stress that may increase vulnerability to dementia we investigated the persistent impacts of combined chronic ethanol exposure and stress on mice when they reached midlife. We exposed young adult mice to excessive alcohol and chronic stress then assessed long-term effects after three months of prolonged abstinence. We assessed cognitive function, including cognitive flexibility, using the Barnes maze and evaluated cellular markers of LC dysfunction, including oxidative stress. Our findings demonstrate that a history of excessive alcohol and stress led to long lasting negative impacts on cognition and LC integrity. These results provide important insights into the mechanisms linking alcohol use and stress with accelerated age-related cognitive decline and may inform future strategies for early diagnosis and prevention of dementia in individuals with AUD.

## Materials and Methods

### Subjects

C57BL/6J mice (n=55), 5 months old at the beginning of the experiment (Strain 664, Jackson Labs, Bar Harbor, ME). Mice were single housed on a 12h light/dark cycle (11 am OFF, 11 PM ON), with ad libitum access to food and water, except when noted below for limited access alcohol consumption. The experimental outline is illustrated in Fig. 1. All procedures were approved by the Institutional Animal Care and Use Committee at the University of Massachusetts at Amherst in accordance with the guidelines described in the US National Institutes of Health Guide for the Care and Use of Laboratory Animals (National Research Council (U.S.), 2011).

**Figure 1.**
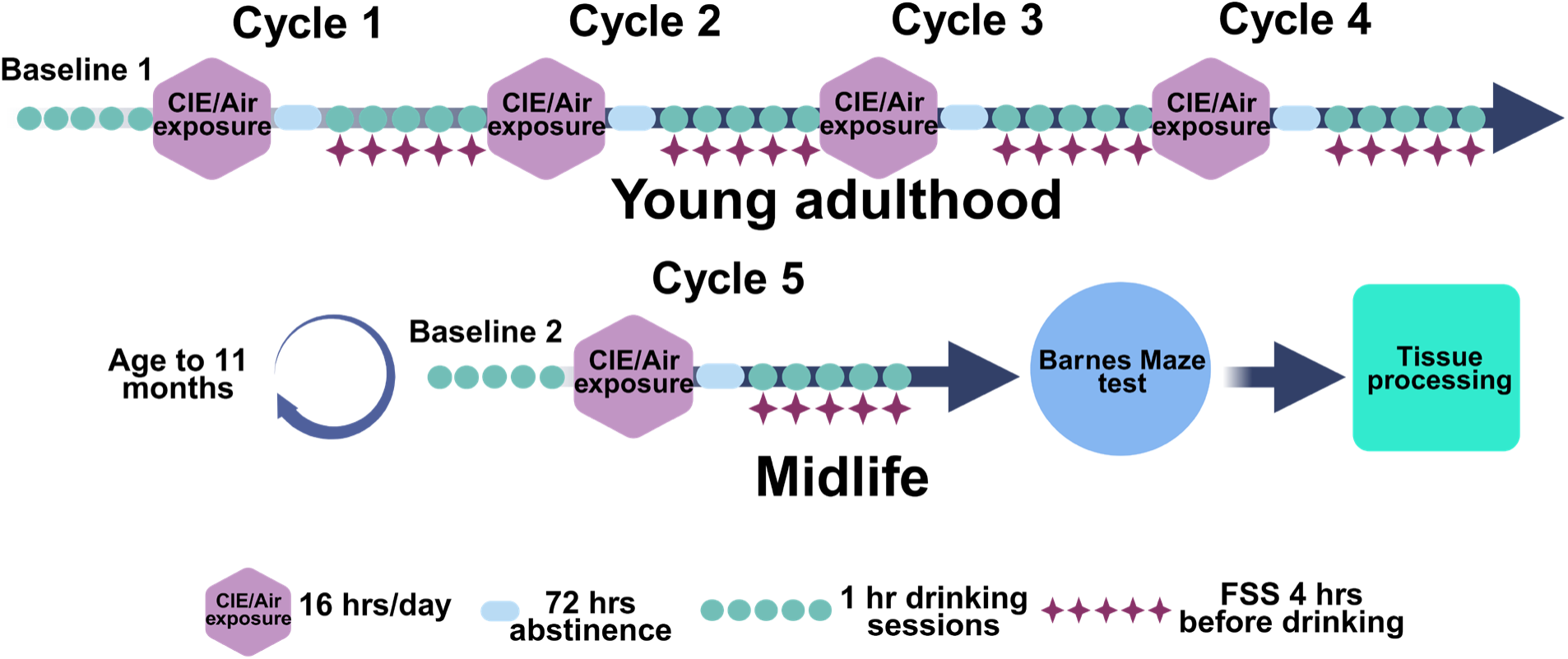
Experimental schedule. Adult mice were given access to ethanol (15%, v/v) 1 hr per day, 5 days/week in their homecage to establish baseline drinking. Subsequently, they underwent four cycles of alternating chronic intermittent ethanol vapor exposure (CIE, 4 days/week, 16 hrs/day,) and repeated daily forces swimm stress (FSS, 5 days/week, 10 mins/day), with control groups exposed to air and no stress. Animals then remained abstinent for 3 months in their homecage. At middle age (11 months), mice repeated baseline drinking, receiving one week access to homecage alcohol (15%, 1hr/day, 5 days/week) followed by a final CIE/FSS to assess stress induced drinking. Following this, they performed a spatial memory based Barnes maze test and reversal. Forty minutes after the last Barnes maze test, brains were collected for RNAscope or immunohistochemical assays to evaluate relevant pathological changes at midlife - 12 months of age.

### Drinking, chronic intermittent ethanol and stress exposure

Mice were given 1 hour limited access daily to 15% ethanol solution in their homecages, Monday to Friday for three weeks, to establish baseline volitional drinking. Animals were weighed weekly to calculate ethanol consumption in g/kg. Mice were classified as either low or high drinkers, using a binary threshold of 2.08 g/kg (males) and 2.23 g/kg (females) average ethanol consumption based on prior work (den Hartog et al., 2020). After stabilization of baseline drinking, mice (n = 27, 15 females, 12 males) underwent cycles of alternating chronic intermittent ethanol (CIE) vapor exposure (1 week) and repeated forced swim stress (FSS) (1 week), with controls (n = 28, 14 females, 14 males) exposed to air and no stress (AIR/NS). Four cycles of CIE and FSS exposure ended at ∼ 8 months of age, at which time mice were returned to their homecages for three months prolonged abstinence and aging out to midlife. At 11 months of age, baseline drinking was tested again before a final cycle of CIE/FSS to evaluate stress induced drinking prior to Barnes maze testing and brain collection at 12 months of age.

During CIE vapor exposure, mice were put in a plexiglass vapor chambers for 16 hours session for 4 consecutive nights, Monday through Thursday. Prior to placement in the chambers, CIE/FSS group received IP loading dose of ethanol (1.6 g/kg), combined with pyrazole as an inhibitor of alcohol dehydrogenase (1 mmol/kg, final volume 20 ml/kg), while the control group received pyrazole only (Becker and Lopez, 2004). On Fridays, blood samples were collected from representative CIE mice to confirm blood ethanol level with alcohol oxidase colorimetric assay (Griffin et al., 2009). CIE chambers kept ethanol levels between 0.17-0.2 mg/dL and produced blood ethanol concentrations (BECs) within the intoxication range (average values (mean ± SEM)) for cycles 1-4: 0.19 ± 0.02 mg/dL, cycle 5: 0.225 ± 0.015 mg/dL). During FSS drinking weeks, mice in the CIE/FSS group were exposed to 10 minutes of FSS in cylindrical tanks of warm tap water (28 °C) while being monitored at the beginning of the dark phase of the light cycle Monday through Friday. Immediately after FSS, mice were gently dried with a paper towel and returned to their homecages. All mice (AIR/NS and CIE/FSS) experienced volitional ethanol drinking sessions (15% ethanol, 1 hour) four hours into the dark phase (Anderson et al., 2016; Lopez et al., 2016; Rodberg et al., 2017).

### Barnes Maze

The Barnes maze is a well-established test used to assess spatial memory and learning as well as cognitive flexibility (Barnes, 1979). Barnes maze apparatus consisted of a circular gray platform (92 cm in diameter) with 20 evenly spaced holes (5 cm in diameter) along the perimeter, mounted on the round table 68.5 cm above the ground (Maze Engineers, Old Orchard Towers, IL). The platform was centrally located within the testing room, with two A4-sized sheets depicting distinct colored signs placed on opposite walls as visual cues for spatial orientation. A black plastic escape box was positioned beneath a designated target hole. Between trials, the platform was thoroughly cleaned with 70% isopropyl alcohol to eliminate residual odors or cues left by previous subjects. To minimize potential spatial learning biases, the target hole position was rotated clockwise every three mice. A white noise generator (∼ 80 dB), combined with bright overhead lighting, encouraged escape behavior. Mice were habituated to the testing room for 30 minutes prior to testing. The Barnes maze protocol consisted of five distinct phases: habituation, training, testing, reversal training, and reversal testing (Attar et al., 2013). Testing of spatial learning and was conducted 48 hours after the final training session. On test day, the escape box was removed, and each mouse was placed inside the start chamber for 15 seconds before being allowed to explore the maze for two minutes. One week following the test session, a reversal learning was conducted with the target hole relocated 180 degrees from its initial position. Mice underwent three training sessions for the new location. The following day, mice underwent a reversal test, during which the escape box was removed, and they were allowed to explore the maze for two minutes. Parameters evaluated included: time spent in the target quadrant, primary latency (the time required to first identify the target), number of errors (nose pokes into incorrect holes before locating the target), distance from the first hole sniffed to target, and search strategy. Additionally, the time spent in the target quadrant, primary latency and error parameters were normalized between animals (as ((value – max value)/(min value – max value))*100 or 1- ((value – max value)/(min value – max value))*100, with higher score being better results), to generate an overall performance score which was the composite sum of these parameters.

### Fluorescence in situ hybridization

For analysis of adrenergic α2A receptors in LC, 8 mice (2 per sex per group) were rapidly anesthetized with isoflurane 40 minutes following the final Barnes maze session. Brains were immediately extracted, flash-frozen on dry ice, and stored at -80°C. Tissue was subsequently sectioned at 12 µm on a Leica CM3050S cryostat (Leica Biosystems, Nußloch, Germany) and thaw mounted on SuperFrost Plus slides (Fisher Scientific, Pittsburg, PA, USA). Fluorescent in situ hybridization (FISH) was performed with the RNAscope Multiplex Fluorescent Assay v2 (Advanced Cell Diagnostics (ACD), Newark, CA, USA) following the manufacturer’s protocol for fresh-frozen tissue (Rodberg et al., 2023). Tissue sections were permeabilized with Protease IV. Probes used were tyrosine hydroxylase *Th* (Mm-Th-C2, 317621-C2, ACD) and *Adra2a* probes (Mm-Adra2a-C3, 425341-C3, ACD). Hybridization was followed by a sequential amplification step using RNAscope Amplifiers 1–3, and HRP signals were obtained by incubating sections with HRP-C2 or HRP-C3 for 15 minutes. Fluorescent signals were developed with Opal fluorophores: Opal 520 (C2 channel, diluted 1:1000 in TSA buffer) and Opal 620 (C3 channel, diluted 1:800 in TSA buffer), each incubated for 30 minutes. Nuclei were counterstained with DAPI for 8 minutes. Finally, slides were mounted using Citi Fluor AF1 Mountant solution (Electron Microscopy Sciences, Hatfield, PA), allowed to dry overnight, and imaged within two weeks post-staining.

### Immunohistochemistry

A subset of mice (n = 16) was perfused with 4% paraformaldehyde (PFA) following the final Barnes maze session. Brain tissue was post-fixed in 4% PFA overnight and subsequently cryoprotected in 20% sucrose azide. Coronal sections (30 µm thick) containing LC were collected. For 4-hydroxynonenal (HNE) immunochemistry, tissue sections were rinsed in Tris-buffered saline (TBS), permeabilized in TBS containing 1% Triton X-100. Sections were blocked for one hour in TBST with 5% normal donkey serum (NDS) and 0.5% bovine serum albumin (BSA), followed by one-hour incubation in donkey anti-mouse IgG Fab fragment (1:33). Sections were then incubated overnight at 4°C with primary antibodies for tyrosine hydroxylase (TH; chicken anti-TH, 1:1000; Aves Lab. Davis, CA) and 4-hydroxynonenal (HNE; mouse anti-HNE, 1:1000), diluted in TBST. Sections were washed in TBST and incubated with fluorophore-conjugated secondary antibodies: donkey anti-chicken Alexa Fluor 488 (1:500) and donkey anti-mouse Alexa Fluor 594 (1:500) for visualization. Control sections were processed without the addition of primary antibodies.

For evaluation of activated caspase-6, tissue sections were washed with TBST (0.2%) and incubated in 1% hydrogen peroxide + 50% methanol for 10 minutes. Tissue was blocked with 3% NDS + TBST for one hour and incubated overnight with rabbit anti-caspase antibody (1:250 dilution in blocking solution). Following the next wash in TBST, tissue was incubated with secondary biotinylated donkey anti-rabbit antibody (1:500). After the next rinse, tissue was incubated in avidin-biotin complex (1:500, ABC Vectastain, Burlingame, CA), with the following chromogenic reaction with DAB-Ni (0.2mg/ml DAB and 6 mg/ml Nickel ammonium sulfate; Fisher Scientific, Pittsburgh, PA). Sections were mounted onto Superfrost Plus slides and dried overnight. The next day, they were counterstained with neutral red for LC visualization, dehydrated and defatted in series of ethanol and xylene, coverslipped with DPX mounting medium.

### Microscopy and image analysis

Image acquisition was done on a Zeiss AxioImager M2 microscope (Zeiss, Munich, Germany). Uniform exposure parameters were applied to ensure consistency. RNAscope puncta analysis was conducted using the open-source software QuPath (v0.5.1, https://qupath.github.io/). The region of interest (ROI) was determined by manually outlining the LC region identified by dense *Th*-positive staining. Individual cells were identified with DAPI using the Cell Detection Tool with the cell expansion parameter set as 8μm. Puncta within the Opal 620 (*Adra2a*) and Opal 520 (*Th*) channels were quantified using the Subcellular Spot Detection Tool. The expression of the α2A-adrenergic receptor was quantified as puncta per cell in *Th*-positive neurons using counts averaged from multiple images per LC. Immunohistochemistry results were quantified using FIJI software (Schindelin et al., 2012). Average HNE fluorescence intensity was quantified within the TH positive LC regions from each subject and compared to background HNE fluorescence intensity from adjacent non-TH regions in the brainstem. Caspase 6 positive cells within the LC, as outlined by neutral red counterstaining, were counted and caspase positive cell density was determined per 100 um^2^ within LC contour bounds.

### Statistical analysis

All statistical analyses were performed on GraphPad Prism Version 10.4.1 (GraphPad Software, La Jolla, CA). Comparisons between groups were made with 2-way RM ANOVA or mixed-effects model with Fisher’s LSD posthoc test where needed, or t-tests (Welch’s t-test) as appropriate. For all analyses, *p* < 0.05 was used as the acceptable α level. Data are presented as mean ± SEM after confirming normality (D’Agostino-Pearson, Shapiro-Wilk, Kolmogorov-Smirnov tests) unless noted otherwise in the figure legend. All graphs were composed using GraphPad Prism.

## Results

### CIE/FSS-induced drinking phenotypes persist in midlife

We and others have previously demonstrated that the combination of CIE/FSS exposure synergistically drives escalation of ethanol consumption in mice compared to either CIE or FSS alone (Anderson et al., 2016; den Hartog et al., 2020). Mice were assigned to one of two experimental groups: a control group receiving only air exposure without stress (AIR/NS) and an experimental group subjected to both CIE and FSS (CIE/FSS). As expected, there was a main effect of CIE/FSS cycles on ethanol consumption (F (1.753, 92.93) = 35.20, P<0.0001; Fig 2A, Two-way RM ANOVA). Animals with a history of CIE/FSS exhibited significantly greater ethanol drinking compared to AIR/NS controls (p < 0.0001, Tukey’s) after four cycles of CIE/FSS (Fig 2A), and maintained high stress induced ethanol consumption compared to AIR/NS at ∼11 months of age (p = 0.0015). Within the AIR/NS or the CIE/FSS group there were no differences in ethanol consumption between 8 and 11 months (p > 0.05, Tukey’s) demonstrating a persistent stress induced drinking phenotype and stable drinking behavior across aging.

Based on prior functional dissociations, mice were classified as either low or high drinkers (den Hartog et al., 2020). As expected, the majority of high drinkers were found in the CIE/FSS group at ∼ 11 months old age (χ², df (13.46, 1), p < 0.0002; Fig 2B)): 82% of Air/NS mice were low drinkers, while 18% were high drinkers. In contrast, 33% of CIE/FSS mice were low drinkers, while 67% were classified as high drinkers. Within both CIE/FSS and Air/NS groups, low and high classifications remained stable across aging in most animals, with 79% of Air/NS mice and 74% of CIE/FSS mice keeping their original categorization (Fig 2C).

**Figure 2.**
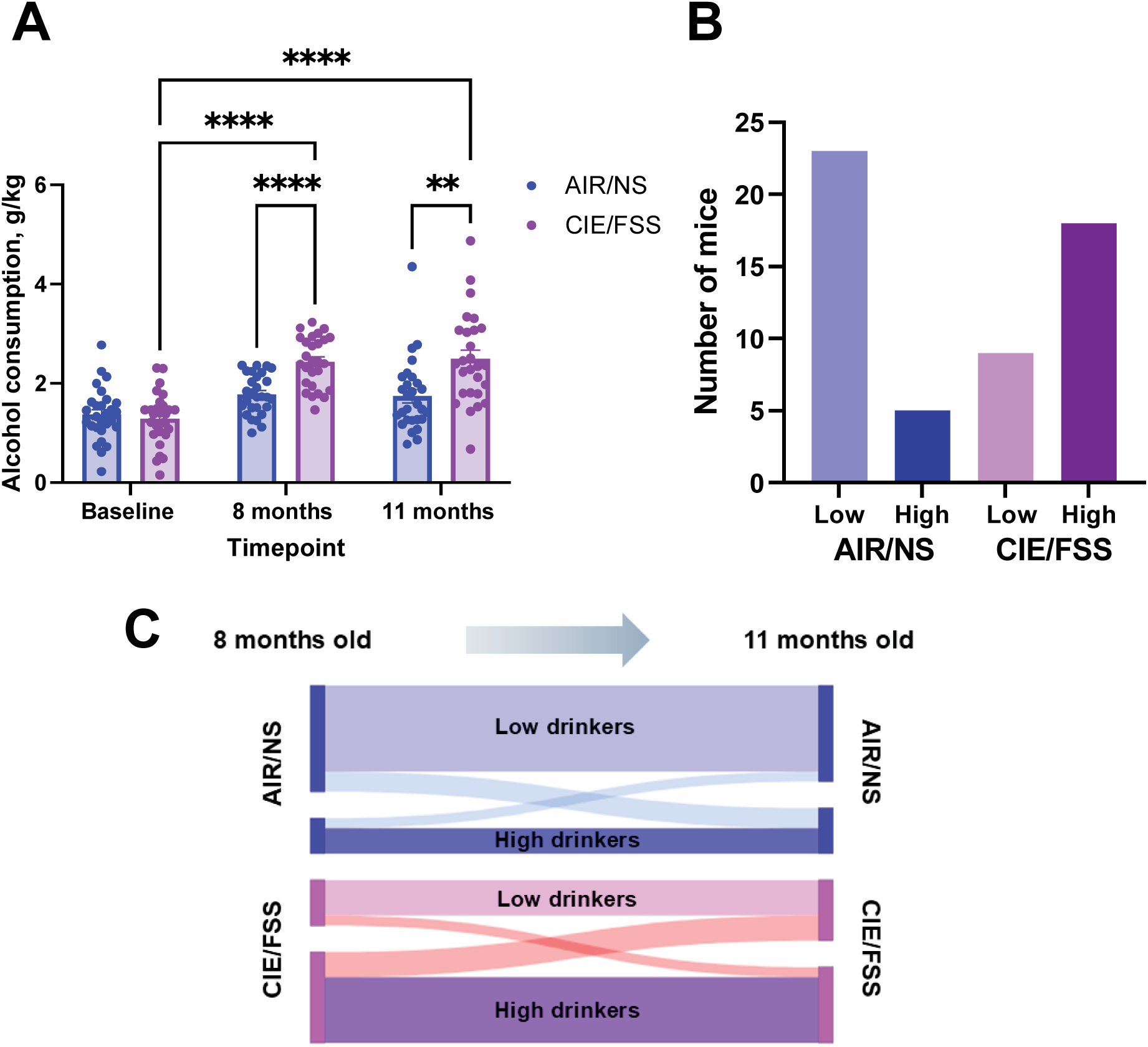
CIE/FSS history creates persistent increases in volitional alcohol consumption. (A) CIE/FSS mice showed an escalation in ethanol consumption that remained in midlife. At midlife AIR/NS group had a majority of low drinkers, while CIE/FSS group had a majority of high drinkers, (C) these phenotypes remained stable after 3 months of abstinence.** p < 0.01; **** p ≤ 0.0001.

### CIE/FSS history disrupted cognitive flexibility, but not spatial learning, in aged mice

The Barnes maze is widely used to assess spatial learning and memory in rodents (Pitts, 2018). In the current study, we included a reversal phase which challenges the animals to adapt to a new escape hole location, giving an opportunity to measure cognitive flexibility (Gawel et al., 2019). As noted above, AIR/NS mice were predominantly low drinkers and CIE/FSS mice were predominantly high-drinkers, thus analysis focused on these two behavioral phenotypes with low power limiting our ability to investigate unique phenotypes in each group (i.e. low drinking CIE/FSS mice). During the initial Barnes test, there were no significant differences in spatial learning between the AIR/NS and CIE/FSS groups in terms of time spent in the target zone, primary latency, or the number of errors (Fig. 3, p>0.05). However, during the reversal phase, which required adaptation to a new target location, differences emerged in cognitive flexibility in the CIE/FSS group (Fig. 3). CIE/FSS mice spent less time in the target zone during the reversal compared to the spatial learning test (Interaction effect of reversal phase x CIE; F (1, 77) = 5.26, p = 0.025; REML ANOVA; Fig. 3A). CIE/FSS mice also made more errors on the reversal phase compared to their prior spatial learning performance (p = 0.048, Fisher’s LSD; Fig 3B). However, there were no significant differences in the primary latency for locating the new escape hole (Fig. 3C). When these individual parameters (target zone time, errors and latency) were integrated as an overall performance score (see Methods), mixed-effect analysis confirmed a significant interaction between reversal performance and CIE/FSS history (F (1, 77) = 4.37, p = 0.04). Post-hoc comparisons showed CIE/FSS had significantly impaired performance during the reversal compared to AIR/NS controls (p = 0.036, Fisher’s LSD) and compared to their own performance on the spatial learning test phase (p = 0.029, Fisher’s LSD).

**Figure 3.**
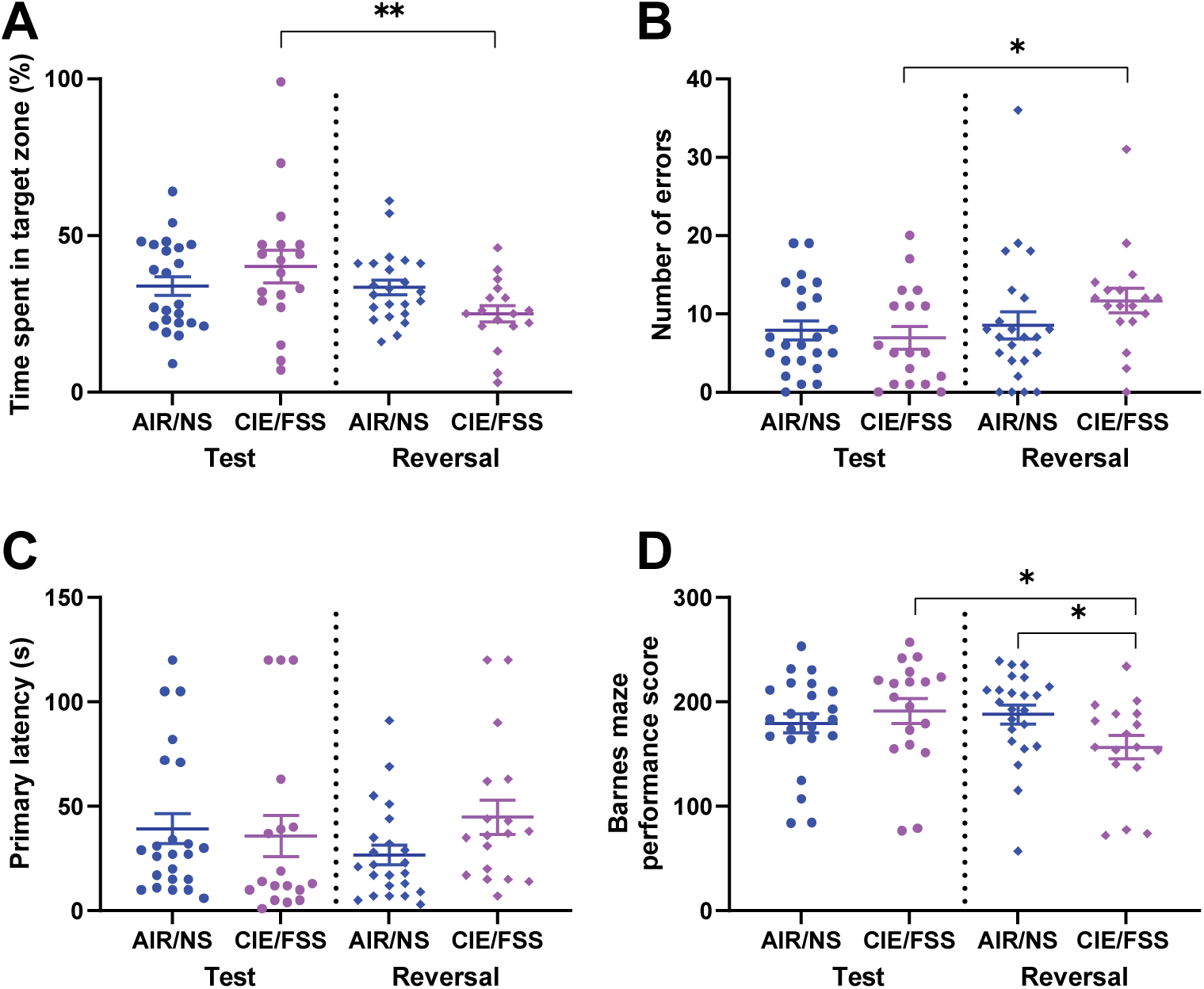
CIE/FSS history leads to impaired flexibility in the Barnes Maze during midlife. (A) CIE/FSS mice reduced time in target quadrant during the reversal test than learning test (p < 0.01); (B) CIE/FSS mice made more errors during the reversal test than learning test (p < 0.05); (C) CIE/FSS mice showed similar primary latency compared to AIR/NS; (D) Collectively, CIE/FSS mice had overall lower reversal performance compared to AIR/NS (p <0.05) and their own test performance (p < 0.05). * indicates p < 0.05; ** p < 0.01.

### CIE/FSS disrupts LC integrity in cognitively impaired aged mice

Oxidative stress is a well-established mechanism of acute alcohol-induced neurotoxicity (Ruiter-Lopez et al., 2025). We wanted to assess whether oxidative stress indicators persist long term after CIE/FSS. We used 4-hydroxynonenal (HNE) as a marker of lipid peroxidation and oxidative damage in LC, a neuromodulatory region critical for cognitive flexibility (Fig. 4A, B). We identified a significant increase in HNE-associated fluorescence in the LC of mice with a history of CIE/FSS compared to AIR/NS (Mann-Whitney test, p < 0.0001, Fig. 4C) at midlife (12 months old) in animals with persistent high drinking and cognitive inflexibility phenotypes, despite 3 months of homecage abstinence. These findings indicate heightened oxidative stress in the LC following combined chronic alcohol and stress exposure, as a mechanism contributing to impaired cognitive flexibility.

To evaluate whether apoptosis was involved in LC dysfunction after CIE/FSS, we examined cleaved (active) caspase-6 expression. Although the CIE/FSS subjects showed a trend toward increased caspase-6 levels in the LC, this effect did not reach statistical significance (p = 0.1 Mann-Whitney test, Fig. 4D) The absence of a robust effect in this case may be attributed to the limited sample size, suggesting that further studies are needed to clarify the impact of alcohol and stress on cell death pathways in LC.

**Figure 4.**
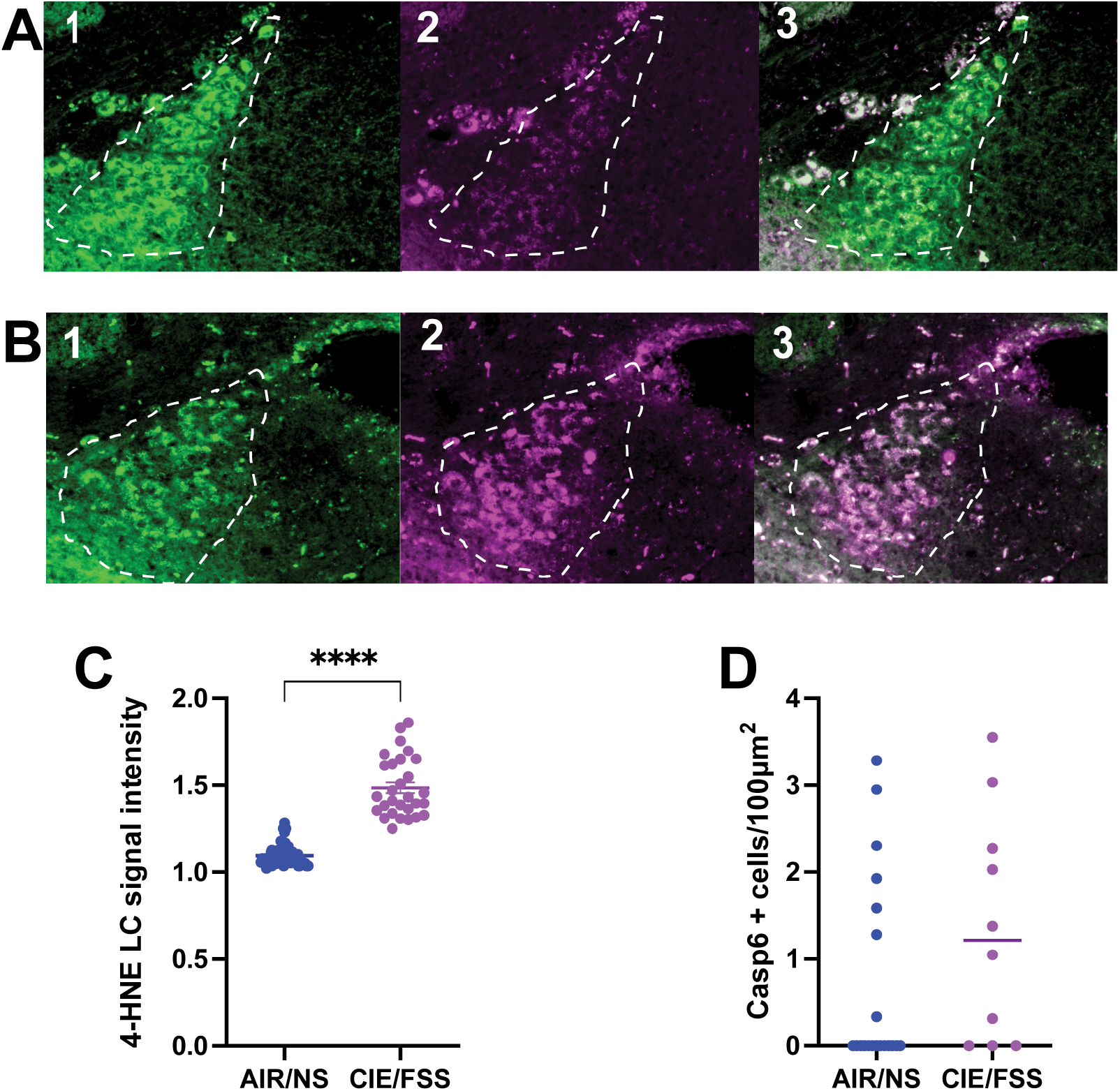
CIE/FSS history increases oxidative stress in LC. (A) Representative images of LC (outlined, TH, green, 1) immunostaining showing oxidative stress (4-HNE, magenta, 2) from AIR/NS and (B) from CIE/FSS subjects with merged composites (3). (C) Quantification of fluorescent HNE intensity relative to background showed pronounced oxidative stress increases in CIE/FSS compared to AIR/NS animals. (D) Activated Caspase-6 density within LC neurons was variable within and between subjects (line at median). **** indicates p < 0.0001.

To investigate whether alcohol and stress disrupt autoregulation of LC function, we assessed α2-AR mRNA expression using RNAscope. Our analysis revealed an 22% reduction in α2-AR expression in LC of CIE/FSS compared to AIR/NS subjects (p = 0.043, unpaired t-test, Fig. 5).

**Figure 5.**
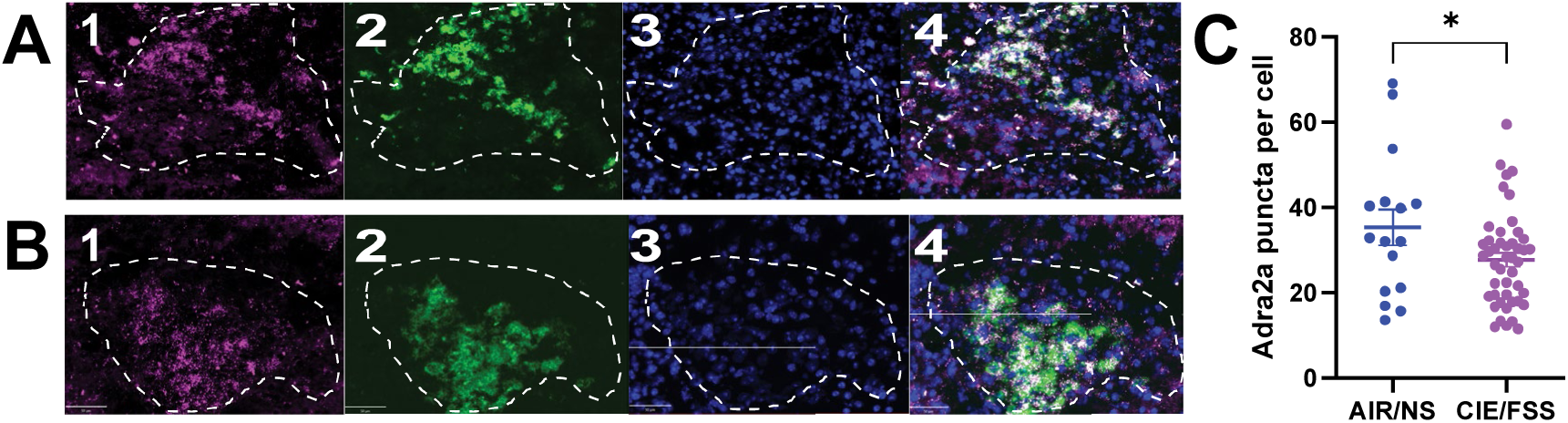
CIE/FSS history decreases Adra2a expression in LC. Representative image of LC (outlined) FISH staining (RNAscope) from an (A) AIR/NS mouse and (B) from a CIE/FSS mouse showing *Adra2a* (magenta, 1) and *Th* (green, 2) probes with DAPI counterstain (blue, 3) and merged composited (4) from. (C) *Adra2a* expression levels were lower in LC cells from mice with a history of CIE/FSS group compared to AIR/NS at 12 months of age. Scale bar = 50um, * indicates P<0.05.

Taken together, these results highlight multiple potential mechanisms by which chronic alcohol and stress exposure may disrupt LC integrity, including oxidative stress, and altered autoreceptor feedback. These effects are indicative of persistent pathological changes in 12-month-old mice, despite primarily receiving alcohol and stress exposure in young adulthood and undergoing extended abstinence. Such pathology could contribute to the cognitive deficits observed in aged animals with a history of heavy alcohol and stress exposure.

## Discussion

The present study demonstrates that the combined action of stress and alcohol at a younger age contributes to persistent heavy drinking, along with cognitive decline and LC pathology in midlife. Using a well-established model, we subjected mice to a chronic alcohol and stress paradigm known to enhance voluntary alcohol consumption (Anderson et al., 2016; Rodberg et al., 2017). The results in our current study demonstrate a lasting impact of chronic alcohol and stress, despite five months of protracted abstinence, increased volitional alcohol intake persisted in animals with a history of chronic alcohol and stress. Furthermore, older mice with a chronic alcohol and stress history showed cognitive flexibility deficits and multiple pathological changes in LC, including increased oxidative stress load and reduced autoregulatory feedback.

CIE/FSS mice demonstrated elevated drinking levels similar to that seen immediately after CIE/FSS exposure despite three months of home cage abstinence. The escalation of alcohol consumption in the CIE/FSS group is consistent with prior evidence linking stress exposure to heightened vulnerability to excessive alcohol use (den Hartog et al., 2020; Rodberg et al., 2017). Notably, this escalated intake was maintained long term, suggesting that alcohol exposure at younger age can induce persistent neuroadaptations in stress- and reward-related neural circuits for long lasting alcohol consumption phenotypes. Therefore, our findings provide further evidence for the long-lasting risk of relapse after a history of stress and alcohol.

In addition to a persistent drinking phenotype, CIE/FSS mice exhibited significant deficits in adapting to new rules in the reversal phase of the Barnes maze during midlife, compared both to the control group and to their own performance in the initial test of spatial memory. These findings align with previous studies from our lab and others, demonstrating detrimental impact of chronic alcohol and/or stress-induced on cognitive flexibility, including reversal learning during, although many of these studies have been evaluated during acute alcohol withdrawal, or early protracted withdrawal (Badanich et al., 2011; Gass et al., 2014; Rodberg et al., 2017; Shnitko et al., 2020), although several studies in humans with AUD have shown long term negative impacts on reversal learning (Martelli et al., 2017)(Kopera et al., 2012). Interestingly, not all aspects of cognition were affected by CIE/FSS. Although reversal learning was impaired, spatial memory, as demonstrated by the initial test in the Barnes maze, was not. These results highlight that a history of prolonged alcohol and stress may preferentially affect frontal cortex-mediated executive functions, such as cognitive flexibility, whereas other hippocampus- and cerebellum-dependent spatial learning and memory processes may be less impacted.

The mechanisms by which chronic alcohol and stress drive cognitive dysfunction remain unknown. There are many allostatic changes and putative targets that may underlie these effects (Nippert et al., 2024). We investigated several potential mechanisms in the LC, a region implicated in stress modulation and critical for cognitive flexibility (Cope et al., 2019; McGaughy et al., 2008; Reyes, 2025). One mechanism with potential to impact cognitive function is oxidative stress in neuronal targets. Alcohol metabolism facilitates the production of reactive oxygen species (ROS), creating imbalance between ROS and antioxidant systems. Alcohol-induced cellular toxicity can result from lipid peroxidation, increased cytokines production, altered metal ion levels, effects on enzyme activity, DNA damage, and more, all of which can lead to cell death and dysfunction (Haorah et al., 2008; Hernandez et al., 2016; Ruiter-Lopez et al., 2025; Wu and Cederbaum, 2003). Thus, alcohol induced oxidative stress may contribute to LC dysfunction and norepinephrine dysregulation of cognitive flexibility. Indeed, immunostaining for 4-HNE, a lipid peroxidation marker, revealed significantly elevated 4-HNE levels in the LC of CIE/FSS mice, providing evidence of persistent increased oxidative stress. These results are consistent with the studies reporting that oxidative stress may also be involved in the etiology of other neuropsychiatric diseases, including dementia, significantly worsening the prognosis and outcomes for patients with concurrent AUD (Marien et al., 2004; Newton et al., 2015; Sayre et al., 1997; Smith et al., 2000).

In addition to oxidative stress, we investigated apoptotic processes in LC. Immunostaining for cleaved caspase-6, a key mediator of programmed cell death(LeBlanc et al., 1999; Theofilas et al., 2018), showed putative increases in caspase-6-positive cells in CIE/FSS mice compared to controls, though the sample size was underpowered for definitive conclusions. Notably, elevated caspase-6 levels have been consistently reported in multiple forms of dementia in LC and other brain regions, suggesting it’s involvement in brain aging and degenerative processes (Guo et al., 2004; LeBlanc et al., 1999; Theofilas et al., 2018; Wang et al., 2015). The potential role for caspase 6 in alcohol induced neural dysfunction warrants further exploration.

Another potential mechanism contributing to alcohol and stress impacts on cognitive function is disrupted autoinhibition in the LC via the α2a-adrenergic receptor. FISH analysis revealed significantly reduced *Adra2A* gene expression in the LC of CIE/FSS mice compared to controls. These findings have important implications as α2 agonists are used clinically to manage acute alcohol withdrawal (Muzyk et al., 2011). Chronic alcohol exposure has been shown to alter expression of other adrenergic receptors in the medial prefrontal cortex and central amygdala, affecting cognitive and emotional processing in conjunction with α2 adrenoreceptors (Athanason et al., 2023). In addition to the consequences of alcohol and stress in the current study, loss of α2a-adrenergic receptors specifically in LC has previously been documented in dementia patients and may be a pathological hallmark (Szot et al., 2006). Further research is needed to clarify the implications of targeting α2a-adrenergic receptor signaling balance in the context of alcohol-related cognitive decline.

Looking forward to future studies, one potential driver of these long-term allostatic changes after chronic alcohol and stress may involve dysregulation of the hypothalamic-pituitary-adrenal (HPA) axis, which plays a crucial role in stress responses. HPA malfunction has been associated with AUD and a higher risk of relapse (Lu and Richardson, 2014; Richardson et al., 2008). LC-NE, in turn, can have an excitatory influence on the HPA axis in response to stress (Reyes, 2025). Disruption of LC function, especially autoinhibition, may further impair HPA axis regulation, exacerbating maladaptive stress responses and contributing to alcohol misuse as a maladaptive coping strategy.

Overall, our findings highlight the long-lasting impact of alcohol and stress exposure on cognitive function, with specific impairments in cognitive flexibility persisting into midlife. These deficits are associated with increased oxidative stress, and adrenergic dysregulation within the LC, a key region mediating cognitive flexibility via its forebrain projections. Although our study provides valuable insights into these mechanisms, several important questions remain. Further characterization of physiological changes in both the LC and frontal cortex will be essential for developing a comprehensive understanding of how these brain regions interact to mediate the long-term consequences of chronic alcohol and stress exposure. The current study provides a foundation for future research into the mechanisms underlying AUD-associated cognitive decline and its relationship to dementia risk. It further highlights potential therapeutic targets in LC, such as oxidative stress pathways and norepinephrine signaling, for managing the long-term consequences of alcohol and stress exposure on cognitive aging.

## Acknowledgements

This work was supported by PHS award U01 AA025481 to DEM and EMV.

The authors have no conflicts of interest to disclose.

## Notes

### Competing Interest Statement

The authors have declared no competing interest.

